# Focal adhesion kinase promotes metastasis in BRAF-mutant melanoma

**DOI:** 10.64898/2026.03.19.712212

**Authors:** Karly A. Stanley, MiKaela N. Field, Adriene M. Pavek, A. Paulina Medellin, Savannah N. Pettey, Mercedes Randhahn, Tursun Turapov, Gennie L. Parkman, David A. Kircher, Benjamin Izar, Alexandra Young, Matthew W. VanBrocklin, Sheri L. Holmen

## Abstract

Despite the availability of several FDA-approved therapies, metastatic melanoma remains a significant clinical challenge, particularly for patients with brain metastases, which frequently represent the site of treatment failure and a major cause of melanoma-related mortality. Melanoma exhibits a strong propensity to metastasize to the brain, yet the molecular mechanisms driving this lethal progression remain incompletely understood, limiting the development of effective treatment options. Building on our prior discovery that focal adhesion kinase (FAK) is a key mediator of AKT1-driven brain metastasis, we sought to validate the role of FAK in melanoma progression and metastatic dissemination. Using complementary autochthonous and syngeneic mouse models of BRAF-mutant melanoma, we evaluated the impact of FAK expression on overall survival, primary tumor growth, and metastasis. Through the generation of targeted FAK mutants, we distinguished kinase-dependent from kinase-independent functions and demonstrate that FAK promotes melanoma metastasis in a kinase-dependent manner. Furthermore, we establish that FAK functions downstream of PTEN to drive metastatic progression. Collectively, these findings support the therapeutic potential of FAK inhibition, either alone or in combination with existing treatments, to more effectively combat metastatic melanoma and inform the development of emerging FAK-targeted therapies.

## Background

Cutaneous melanoma is one of the most lethal forms of skin cancer due to its high rates of metastasis. While there are several FDA-approved therapies for advanced melanoma, including immunotherapies and targeted therapies, ∼50% of patients do not respond to current treatments^1–3^. Among those who do respond, many eventually develop resistance, and targeted therapies are only approved for patients with melanoma harboring mutant BRAF. As a result, not all patients benefit, and the majority of melanoma-related deaths result from distant metastases^4^. Therefore, it is critical to further our understanding of the mechanisms that drive melanoma metastasis such that more effective therapeutic strategies can be developed to improve outcomes in these patients. We and others have demonstrated that activation of the phosphoinositide 3-kinase (PI3K)/protein kinase B (AKT) signaling pathway promotes melanoma metastasis. Specifically, we discovered that activation of AKT1 signaling drives melanoma metastasis in an autochthonous mouse model of cutaneous melanoma^5,6^. These data implicate PI3K/AKT pathway inhibition as a possible treatment strategy for melanoma; however pharmacologic inhibitors of this pathway have shown limited efficacy and/or intolerable toxicity^7^. To identify alternative targets, we utilized a proteomics approach to identify downstream effectors of AKT1 signaling that promote the brain metastatic phenotype. These studies revealed a significant increase in the levels of phosphorylated focal adhesion kinase (FAK) in brain metastatic compared to non-brain metastatic melanomas^6^.

FAK is a non-receptor tyrosine kinase that canonically signals downstream of integrins and growth factors to regulate biological functions at focal adhesions including cell growth, migration, and invasion. FAK consists of three functional domains: an N-terminal 4.1 ezrin, radixin, moesin homology (FERM) domain, a kinase domain, and a C-terminal focal adhesion targeting (FAT) domain. The FERM domain is a highly conserved domain commonly found in cytoskeletal-associated proteins and facilitates protein localization to the cell membrane. The primary binding partners of the FERM domain include transmembrane receptors, integrins, and phosphoinositides^8^. The FERM domain also regulates FAK signaling through autoinhibitory interactions with the kinase domain, blocking FAK autophosphorylation at Y397 and preventing full FAK activation^9–11^. This autoinhibitory interaction is relieved upon conformational changes stimulated by lipid binding, extracellular matrix binding, and integrin signaling. Autophosphorylation of Y397 creates a Src homology 2 (SH2)-domain binding site, facilitating Src recruitment and phosphorylation of FAK residues Y576 and Y577 within the activation loop of the kinase domain, further activating FAK. Src also phosphorylates Y861 and Y925, which facilitates interactions with downstream signaling partners such as p130Cas-Crk and Grb2, respectively, contributing to adhesion dynamics, cell migration, and survival signaling^9–14^. Finally, the FAT domain is responsible for localization to focal adhesions through direct binding with other focal adhesion proteins including talin and paxillin^9,10,12^.

In addition to its canonical kinase-dependent roles at focal adhesions, FAK exerts a variety of kinase-independent functions, primarily through its scaffolding activity. For example, FAK has been shown to translocate to the nucleus, where it functions as a scaffold to regulate the transcription of genes like *IGFBP3* or to facilitate MDM2-mediated ubiquitination of p53^15,16^. Gene amplification of *PTK2*, which encodes FAK, has been observed in various cancers including breast cancer, ovarian cancer, and glioma, where FAK expression correlates with worse patient outcomes^12,13,17–21^. Despite the well-known role of FAK in cell migration and invasion, and the association of FAK expression with poor patient outcomes, the role of FAK in promoting melanoma metastasis has been largely unexplored. Based on our previous findings and the pro-tumorigenic roles of FAK reported in the literature^22,23^, we sought to understand the role of FAK kinase activity on melanoma tumor progression and metastasis.

In this study, we first investigated whether *PTK2* expression predicts outcomes in human melanoma. Our analysis revealed a strong link between high *PTK2* expression and worse overall survival. In patients with BRAF-mutant melanoma treated with standard of care dabrafenib plus trametinib (BRAF/MEK inhibitors), high FAK expression was associated with significantly reduced treatment efficacy indicating that elevated FAK signaling may confer intrinsic resistance to targeted therapy in BRAF-mutant driven disease. These results suggest that FAK is a key contributor to melanoma aggressiveness and treatment resistance. To elucidate the role of FAK in driving melanoma metastasis, we generated a panel of FAK mutants designed to disrupt specific functional protein domains. We found that FAK expression drives melanoma metastasis and that this phenotype is dependent on FAK kinase activity. Further, we discovered that the FERM domain is not required to promote metastasis. Additionally, we investigated the interplay of FAK signaling along the PI3K/AKT signaling axis and found that FAK functions downstream of PTEN to drive melanoma metastasis. These results suggest that high FAK expression is a poor prognostic indicator for melanoma patient survival, validate kinase-active FAK as a driver of melanoma metastasis downstream of PTEN and AKT1, and support the proposed use of ATP-competitive FAK inhibitors for the prevention and/or treatment of metastatic melanoma.

## Methods

### Mice and genotyping

All animal experimentation was performed in AAALAC approved facilities at the University of Utah. All animal protocols were reviewed and approved prior to experimentation by the Institutional Animal Care and Use Committee (IACUC) at the University of Utah. *Dct:TVA;Braf^CA^;Cdkna2a^lox/lox^;Pten^lox/lox^* mice were maintained on a mixed C57BL/6 and FVB/N background by random interbreeding. DNA from tail biopsies was used to genotype for the TVA transgene, *Braf^CA^*, *Cdkn2a^lox/lox^*, *Pten^lox/lox^,* and wild-type alleles as described^24^. Both male and female newborn through adult mice were used in this study. Mice were fed a combination of Teklad Global 2920X and Teklad 3980X (Inotiv), supplemented with DietGel 76A and HydroGel (Clear H_2_O) post-weaning. One-week post weaning, mice were transitioned to Teklad Global 2920X.

### Viral constructs and propagation

Avian retroviral vectors used in this study are replication-competent Avian Leukosis Virus splice acceptor and Bryan polymerase-containing vectors of envelope subgroup A [designated RCASBP(A) and abbreviated RCAS]. A Gateway compatible RCAS destination vector has been described^25^. The RCAS-Cre, pCR8 AKT1^E17K^ entry vector, and RCAS-AKT1^E17K^ constructs have also been described ^6^. Mouse melanocyte cDNA and FAK-specific primers were used to amplify FAK via PCR. The product was TOPO cloned into the gateway compatible pCR8 TOPO vector (Thermo Fisher, Waltham, MA), then subcloned into the RCAS destination vector. RCAS-FAK was used as a template to generate the FAK mutant constructs. All mutants were engineered with an N-terminal HA-epitope tag. RCAS-FAK was used as a template to generate the Y397E mutant construct using the Aligent QuikChange Site-Directed Mutagenesis Kit (Agilent Technologies, Santa Clara, CA) with mutant specific overlapping primers. All viral vectors were verified via DNA sequencing. Primer sequences are available upon request. Viral infection was initiated by polyethyleneimine (PEI) transient transfection of proviral DNA into sub-confluent DF-1 avian fibroblast cells. Successful production of the virus was confirmed by immunoblot for the avian leukosis viral capsid p27 [Immunology consultants laboratory (ICL) clone #C62. Catalog #MALV-30A-6C2], HA, and the protein of interest. Gateway cloning was used to recombine the pCR8 AKT1^E17K^, pCR8 FAK-WT, pCR8 FAK^K454R^, pCR8 FAK^NDEL^, or pCR8 FAK^Y397E^ entry clones with the pDEST FG12-Luciferase-IRES-EGFP lentiviral destination vector (Gene Universal) to generate lentiviral vectors containing each of these genes.

### Generation of cell lines expressing wild-type and mutant FAK

A *Pten*-deficient isogenic variant of the YUMM3.2 parental cell line, a gift from Marcus Bosenberg, was generated via CRISPR/CAS9 as previously described^26^. To generate virus from the lentiviral vectors described above, psPAX2, and pCMV-VSV-G plasmids were co-transfected into HEK 293FT cells (Thermo Fisher; R70007) using lipofectamine 3000 (Thermo Fisher). Viral supernatant produced from the transfected cells was used to infect YUMM3.2 Pten^−/−^ cells. Expression of the exogenous proteins was confirmed by immunoblot for HA.

### Cell culture

DF-1 cells were grown in DMEM-high glucose media (Thermo Fisher) supplemented with 10% FBS (Atlas Biologicals) and 0.5 µg/mL gentamicin (Thermo Fisher) and maintained at 39°C in 5% CO_2_. YUMM3.2 cells were grown in F12/DMEM media (Thermo Fisher) supplemented with 10% FBS, 1 µg/mL penicillin-streptomycin (Thermo Fisher) and 1 µg/mL non-essential amino acids (Thermo Fisher) and maintained at 37°C in 5% CO_2_.

### Proliferation and transwell migration assays

To measure cell confluence as a read out for cell proliferation, all YUMM3.2 cell lines were plated in triplicate in a 96-well plate at 3,000 cells per well. Confluence was assessed over time at 37°C using an IncuCyte® Zoom Live Cell Imaging instrument with data analyzed using IncuCyte® Analysis Software (Sartorius) at four-hour intervals. To measure wound healing as a readout for migration, 150,000 cells from each YUMM3.2 cell line were plated in triplicate in a 96-well plate and supplemented with F12/DMEM media supplemented with 10% FBS (Atlas Biologicals), 1 ug/mL penicillin-streptomycin (Thermo Fisher), and 1 ug/mL non-essential amino acids (Thermo Fisher). The Sartorius 96-well Woundmaker tool was utilized to create a consistently sized wound in all wells simultaneously. Wound closure was monitored over 72 hours using the Incucyte S3 live cell imaging system. Transwell migration assays were also performed to analyze cell migration. In this assay, all YUMM 3.2 cell lines were plated in F12/DMEM serum-free media (Thermo Fisher) supplemented with 1% BSA (Cell Signaling Technology, Danvers, MA), 1 ug/mL penicillin-streptomycin (Thermo Fisher), and maintained at 37°C. After 24 hours, 30,000 serum-starved cells were re-suspended in F12/DMEM serum-free media (Themo Fisher) supplemented with 1% BSA (Cell Signaling Technology), 1 ug/mL non-essential amino acids (Thermo Fisher) and plated on the surface of each transwell (6.5 mm, 8 um pores) (Corning). As a chemoattractant, 800 uL of F12/DMEM media supplemented with 10% FBS (Atlas Biologicals), 1 ug/mL penicillin-streptomycin (Thermo Fisher), and 1 ug/mL non-essential amino acids (Thermo Fisher) was added to the bottom of the chamber. Media on the apical surface of the transwells was refreshed daily. On day 5, transwells were rinsed in 1 × PBS, incubated in 100% methanol for 100 minutes, and stained with crystal violet for 10 minutes. Cells on the apical surface of the transwells were removed using a cotton swab. Cells on the basolateral surface of each well were imaged, then incubated in 20% acetic acid in H_2_O for 10 minutes on a rotating orbital to dissolve the crystal violet. Quantification was performed in ImageJ by thresholding and particle analysis, and values represent the mean migrated cells per field normalized to control.

### In vivo viral infections and RCAS tumorigenesis studies

Infected DF-1 cells from a confluent culture in a 10 cm dish were trypsinized, pelleted, resuspended in 100 µL of Hank’s Balanced Salt Solution (HBSS) (Thermo Fisher), and placed on ice. Newborn mice were injected subcutaneously near the ear pinnae with 50 µL of suspended RCAS-AKT1^E17K^, RCAS-FAK^WT^, RCAS-FAK^NDEL^, or RCAS-FAK^Y397E^ cells mixed with RCAS-Cre cells. Upon weaning, mice were monitored daily for tumor growth. Tumor burden was quantified three times weekly using digital caliper measurements using the following formula to calculate tumor volume: (Length x Width^2^)/2.

### In vivo syngeneic allografts

Six- to ten-week-old C57BL/6-Tg(CAG-luc-EGFP)GH (glowing head) mice were injected subcutaneously into the dorsal area near the scapula with 2 × 10^5^ YUMM3.2 Pten^−/−^ cells co-expressing AKT1^E17K^, FAK^WT^, FAK^K454R^, FAK^NDEL^, or FAK^Y397E^ and Luciferase-IRES-EGFP. Mice were monitored daily for tumor formation and tumors were visualized and measured weekly using bioluminescent imaging (BLI). Tumor burden was quantified using bioluminescent radiance values (photos/second/cm^2^/sr) and digital caliper measurements.

### Bioluminescent imaging

Mice were injected intraperitoneally with 100 µL of 16.7 mg/mL D-Luciferin 10 minutes prior to image acquisition. The IVIS Spectrum (Xenogen, Alameda, CA) was used to acquire images at one-week intervals beginning one-week post-injection and continued until euthanasia. Mice were euthanized in accordance with IACUC-approved humane endpoints, including tumor burden, signs of distress or declining health, or at the predetermined experimental endpoint. At sacrifice, the primary tumor, lungs, liver, and brain were imaged *ex vivo* to confirm luciferase expression and to detect metastases. Living Image software (version 4.5.2) was used to compile images.

### Histology and histochemical staining

Euthanized mice at their experimental endpoints and subjected to a full necropsy. Brain, lung, liver, and primary tumor tissues were fixed in formalin overnight, dehydrated in 70% ethyl alcohol, and paraffin embedded. Sections (4 µm) were adhered to glass slides and stained with hematoxylin and eosin (H&E) by the Biorepository and Molecular Pathology Shared Resource at the University of Utah.

### Immunoblotting

Cell lysates were suspended in 100 mM Tris-HCl, 4% SDS, 20% glycerol, and 10% DTT (added fresh). All lysates were incubated at 95°C for 10 minutes, separated on an 8%-16% Tris-glycine polyacrylamide gel (Thermo Fisher), and transferred to nitrocellulose membranes for immunoblotting. Membranes were incubated in blocking solution composed of 0.1% Tween-20 in 1 × TBS (TBS-T) with 5% BSA (Cell Signaling Technology) or 10% non-fat dry milk (NFDM) for 1 hour at room temperature. Blots were incubated in the primary antibody at 4°C overnight with constant shaking and washed in TBS-T. Blots were then incubated in anti-mouse or anti-rabbit IgG-HRP secondary antibody diluted 1:1,000 in TBS-T for 1 hour at room temperature with constant shaking and washed in TBS-T. Enhanced chemiluminescence (ECL, Amersham) was used according to the manufacturer’s specifications and blots were imaged using the Azure Imaging system (Azure Biosystems).

### Survival analysis and PTK2 mRNA expression from patient samples

Samples were analyzed from the Caris CODEai™ data platform. Patient molecular cohorts were selected for analysis based on status of melanoma, and high *PTK2* expression [defined as ≥ 75^th^ percentile TPM in melanoma cases (264.37 tpm)], or low *PTK2* expression [defined as < 25^th^ percentile TPM in melanoma cases (105.57 tpm)]. Real-world overall survival (OS) information was obtained from insurance claims data. OS was calculated from time of diagnosis or from start of treatment (with BRAF/MEK inhibitors dabrafenib or trametinib) until last contact. Patients without contact/claims data for a period of at least 100 days were presumed deceased. Conversely, patients with a documented clinical activity within 100 days prior to the latest data update were censored in the analysis. Kaplan-Meier estimates were calculated for molecularly defined patient cohorts, and significance was determined as *p* < 0.05.

## Results

### High FAK Expression Correlates with Worse Overall Survival in Cutaneous Melanoma

Based on previous results from our mouse models, which demonstrated elevated levels of phosphorylated FAK Y397 in brain metastatic melanomas^6^, we first sought to determine whether FAK expression in human melanomas was predictive of poor prognosis. Using the Caris CODEai™ clinico-genomic database, we analyzed a dataset consisting of over 13,000 melanoma patient samples and separated them into quartiles based on mRNA expression of *PTK2*, the gene that encodes the FAK protein. The diagram of each cohort and the *PTK2* expression distribution are shown in Supplemental Figure 1, and additional characteristics of each cohort are included in Supplemental Table 1. High *PTK2* mRNA expression in melanoma [defined as ≥ 75^th^ percentile transcripts per million (TPM)], is significantly associated with worse overall survival (OS) compared with patients whose tumors exhibit low *PTK2* mRNA expression (defined as < 25^th^ percentile TPM). Patients with melanoma exhibiting high *PTK2* expression had a median OS of 78.368 months [95% confidence Interval (CI): 73.433 months (m) – 84.06 m] compared to 91.725 m (95% CI: 86.066 m – 99.588 m) in patients with *PTK2*-low expressing melanoma, which corresponds to a median difference in OS of 13.357 m and a hazard ratio (HR) of 0.818 (95% CI: 0.742 – 0.902; *P* < 0.0001) (Figure 1A). These data suggest that elevated *PTK2/*FAK expression may serve as a negative prognostic biomarker in melanoma. The strong association between high melanoma *PTK2* expression and poor survival supports the hypothesis that FAK plays a functional role in melanoma progression and possibly resistance to therapy.

**Figure 1:**
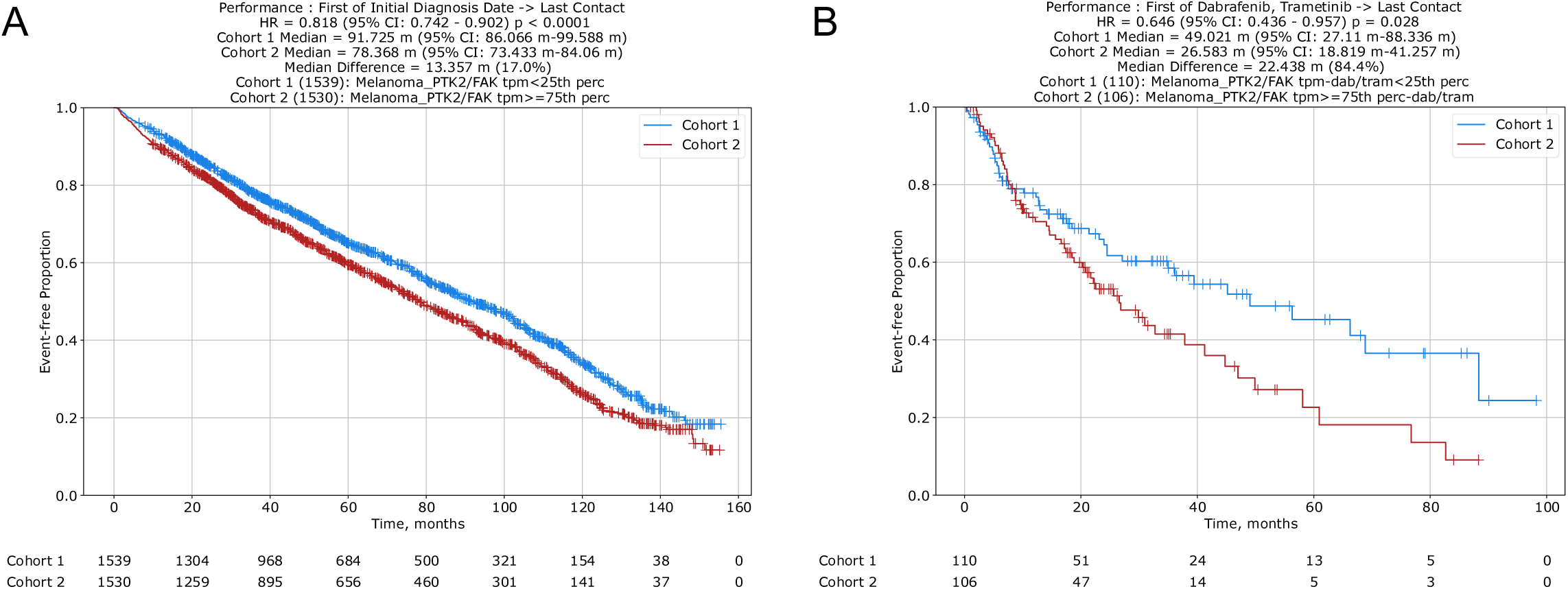
High FAK expression observed in human melanoma samples correlates with worse prognosis. Kaplan-Meier survival analysis from a clinico-genomic database, comparing low (Cohort 1) and high (Cohort 2) *PTK2* (FAK) expression in melanoma to evaluate (A) overall survival using a performance metric of initial diagnosis date to last contact and (B) overall survival using a performance metric of treatment with dabrafenib and trametinib in BRAF-mutant melanoma samples to last contact. High *PTK2* (FAK) expression is defined as ≥ 75^th^ percentile transcripts per million (TPM) in melanoma cases and low *PTK2* (FAK) expression is defined as < 25^th^ percentile TPM in melanoma cases.

To test whether *PTK2* expression influences the response to targeted therapy, we evaluated a cohort of patients with mutant BRAF-driven melanoma treated with the standard of care BRAF and MEK inhibitor combination of dabrafenib and trametinib. High melanoma *PTK2* mRNA expression (≥75th percentile) was associated with significantly shorter overall survival compared to patients with low melanoma *PTK2* expression (<25th percentile); the high *PTK2* cohort had a median OS of 26.583 m (95% CI: 18.819 m – 41.257 m) compared to 49.021 m (95% CI: 27.11 m – 88.336 m) in those with low melanoma *PTK2* expression (below the 25th percentile), corresponding to a median difference in OS of 22.438 m and a HR of 0.646 (95% CI 0.436 – 0.957; *P* = 0.028) (Figure 1B). The reduced efficacy of dabrafenib plus trametinib in the high FAK expression cohort suggests that elevated FAK signaling may confer intrinsic resistance to targeted therapy in the context of mutant BRAF-driven melanoma, in agreement with others^27^. These findings support the role of FAK not only as a negative prognostic biomarker, but also as a potential contributor to therapeutic resistance.

### FAK Kinase Activity Promotes Melanoma Cell Migration and Invasion, Independent of Proliferation, In Vitro

To elucidate the multifaceted roles of FAK in melanoma metastasis, a series of FAK mutants were engineered to modulate its function. Two mutants were designed to enhance FAK kinase activity: one consisting of a 375-amino-acid N-terminal deletion (FAK^NDEL^), which disrupts the auto-inhibitory interactions between the FERM and kinase domains, and a second mutant that incorporates a phospho-mimetic tyrosine-to-glutamic acid substitution at residue Y397 (FAK^Y397E^), the primary FAK autophosphorylation site. The deletion of the N-terminal FERM domain has previously been reported to enhance FAK activity^28,29^ and the Y397E mutation has been demonstrated in the literature as a phospho-mimetic mutant that also enhances FAK activity^30,31^. Additionally, a kinase-deficient mutant (FAK^K454R^)^32^ was generated to delineate kinase-independent functions and explore the scaffolding capabilities of FAK. These constructs, along with wild-type (wt) FAK (FAK^WT^), were integrated into lentiviral vectors co-expressing luciferase and green fluorescent protein (GFP). Virus was produced and used to infect Yale University Mouse Melanoma 3.2 (YUMM3.2) cells, which harbor BRAF^V600E^ and are deficient for *Cdkn2a*. These cells were further modified using CRIPSR/Cas9 gene editing to delete *Pten*. Each FAK cDNA was tagged with an N-terminal HA epitope to distinguish exogenous from endogenous FAK expression. A physiologically relevant active AKT1 mutant, AKT1^E17K^, was utilized as a positive control due to its ability to promote melanoma brain metastasis *in vivo*^5,6^. Immunoblot analysis for the HA epitope tag confirmed the expression of AKT1^E17K^ and all FAK constructs in YUMM3.2 cells (Figure 2). The activity of each protein was also evaluated by immunoblot. The level of P-FAK at Y397 was reduced in the cells expressing the kinase-dead FAK^K454R^ mutant as expected.

**Figure 2:**
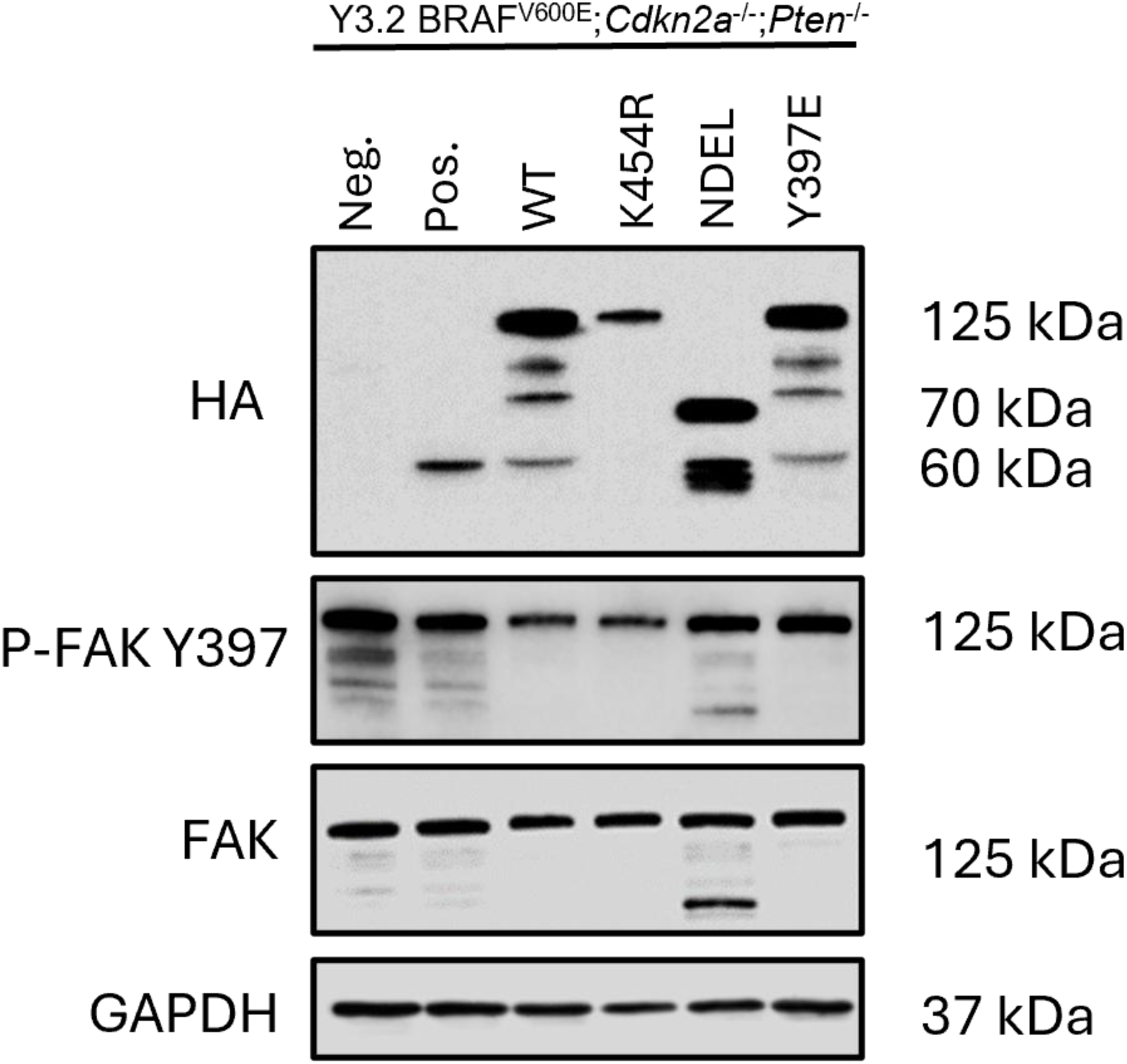
Validation of exogenous FAK construct expression in YUMM3.2;Pten^−/−^ cell lines. Expression of AKT1^E17K^, FAK^WT^, active FAK mutants (FAK^NDEL^ and FAK^Y397E^), and kinase-dead (FAK^K454R^) was confirmed by anti-HA immunoblot. Phosphorylated FAK (Y397) as well as total FAK was assessed as a measure of FAK activity. GAPDH was included as a loading control.

Upon confirmation of exogenous protein expression, we examined the effects of FAK activity on cell confluence as a measure of cell proliferation. Consistent with prior studies, the expression of AKT1^E17K^ did not significantly alter cell proliferation^6^. Similarly, with the exception of FAK^NDEL^, expression of the FAK constructs did not significantly alter cell proliferation, and cells expressing FAK^NDEL^ proliferated slower than the negative control (Figure 3A). These data suggest that active FAK does not confer a proliferative advantage in this context. To evaluate the effects of FAK on cell migration, a wound healing assay was performed whereby a uniform scratch was created using the Sartorius 96-well Woundmaker Tool. Closure of the wound was monitored using the Incucyte S3 live cell imaging system. Cell lines expressing constitutively active AKT1^E17K^ or the hyperactive FAK^NDEL^ exhibited significantly accelerated wound closure compared to the parental cell line (Figure 3B).

**Figure 3:**
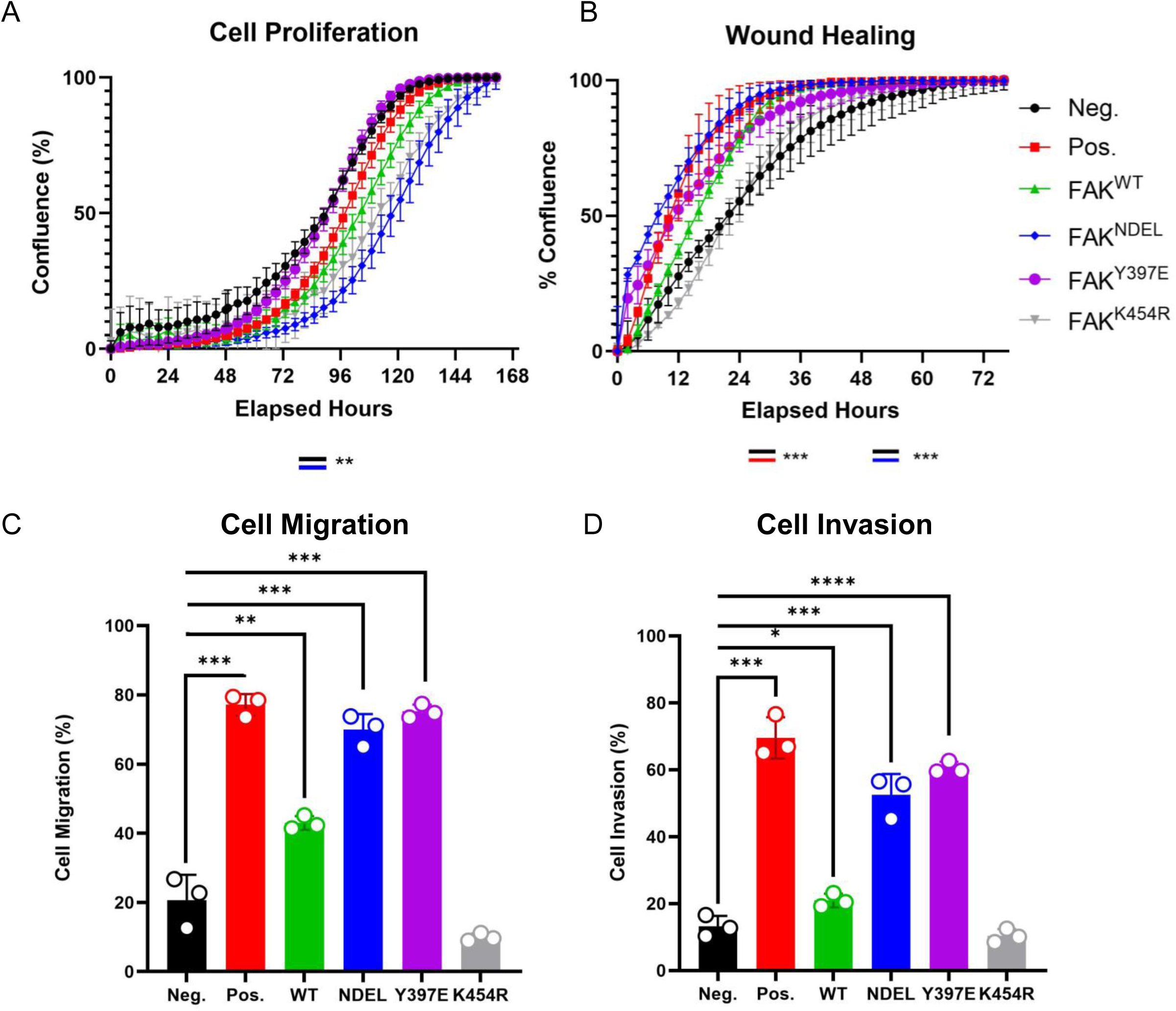
FAK activity in YUMM3.2 cells does not enhance proliferation but promotes migration and invasion *in vitro*. (A) Cell proliferation of YUMM3.2 BRAF^V600E^*;Cdkn2a^−/−^;Pten^−/−^* cells (black line; negative control) engineered to express AKT1^E17K^ (red line; positive control), FAK^WT^ (green line), FAK^NDEL^ (blue line), FAK^Y397E^ (purple line), or FAK^K454R^ (gray line). Cells were plated at 3,000 cells per well in a 96-well plate and cell growth was monitored for one week using the Incucyte S3 live cell imaging system. Cells expressing AKT1^E17K^, FAK^WT^, FAK^Y397E^, or FAK^K454R^ did not significantly alter cell proliferation. In contrast, expression of FAK^NDEL^ slowed proliferation compared with the parental cells. (B) YUMM3.2 control, AKT1^E17K^, FAK^WT^, FAK^NDEL^, FAK^Y397E^, or FAK^K454R^-expressing cells were plated at 150,000 cells per well in a 96-well plate. Once attached, a wound was created in the center of each well simultaneously utilizing the Sartorius 96-well Woundmaker tool. Wound healing was monitored for 72 hours using the Incucyte S3 Live Imaging system. YUMM3.2 cells expressing AKT1^E17K^ or FAK^NDEL^ significantly enhanced wound closure compared to the negative control. (C) YUMM3.2 control, AKT1^E17K^, FAK^WT^, FAK^NDEL^, FAK^Y397E^, or FAK^K454R^-expressing cells were plated on the apical surface of transwells. The ability of cells to migrate through pores towards a chemoattractant. Migrated cells on the basolateral surface were fixed, stained with crystal violet, and imaged. Quantification was performed in ImageJ by thresholding and particle analysis, and values represent the mean migrated cells per field normalized to control. YUMM3.2 cells expressing AKT1^E17K^, FAK^WT^, FAK^NDEL^, FAK^Y397E^ or FAK^Y397E^ enhanced migration compared to the parental cells. In contrast, no significant difference was observed between the parental cells and those expressing FAK^K454^ (D) YUMM3.2 control, AKT1^E17K^, FAK^WT^, FAK^NDEL^, or FAK^K454R^-expressing were plated on the apical surface of transwells above a layer of Matrigel. The ability of cells to invade through Matrigel and pores towards a chemoattractant was measured on day 5. Cells on the basolateral side of transwells were processed as per the migration assay in (C). YUMM3.2 AKT1^E17K^, FAK^WT^, FAK^NDEL^, and FAK^Y397E^-expressing cells significantly increased invasion as compared to control, while FAK^K454R^-expressing cells was not able to promote invasion (p=0.2599). P values are as follows: p < 0.05 (*), p < 0.01 (**), p < 0.001 (***), p < 0.0001.

These findings were corroborated using a transwell migration assay, whereby cells were plated on the apical surface of a transwell chamber in serum-starved media. Full-serum media was added to the bottom of the chamber to promote cell migration through the creation of a chemotactic gradient. After five days, cells that had successfully migrated through the pores of the transwell were fixed and stained in crystal violet for quantification. Cell lines expressing constitutively active AKT1 (AKT1^E17K^), hyperactive FAK (FAK^NDEL^ or FAK^Y397E^), or over-expressing wt FAK (FAK^WT^), exhibited significantly accelerated migration compared to the parental cell line (Figure 3C). In contrast, the kinase-deficient FAK^K454R^ cell line displayed migration rates comparable to the parental cells. To evaluate the effects of FAK activity on cell invasion, the same cell lines were plated into Matrigel-coated transwell chambers. Consistent with our prior observations^6^, expression of AKT1^E17K^ significantly enhanced the ability of cells to invade through the Matrigel (69.49 % +/− 3.57 %) compared with the parental cells (13.20 % +/− 1.82 %; *P* = 0.0001). Whereas FAK^WT^ enhanced invasion 1.5-fold (20.93 % +/− 1.17 %), FAK^NDEL^ enhanced invasion 4.0-fold (52.49 % +/− 3.61 %) and FAK^Y397E^ enhanced invasion 4.6-fold (60.58 % +/− 1.03 %), similar to the 5.3-fold increase in invasion in cells expressing active AKT1 (*P* = 0.0002). In contrast, kinase-dead FAK^K454R^ was unable to promote invasion (10.36 % +/− 1.17 %) demonstrating that kinase activity is required for this function (Figure 3D).

### Loss of FAK Kinase Activity Prolongs Survival and Slows Tumor Growth in a Syngeneic Melanoma Mouse Model

To investigate the role of FAK in melanoma tumor progression and evaluate the impact of FAK kinase deficiency *in vivo*, a syngeneic melanoma model was employed using YUMM3.2 cells expressing the aforementioned AKT1 and FAK mutants alongside EGFP and luciferase, enabling tumor growth tracking via bioluminescent imaging (BLI). These cells, which are syngeneic to C57BL/6 mice, were subcutaneously injected into 6- to 10-week-old C57BL/6 glowing head (GH) mice, which are tolerized to luciferase and EGFP^33^. Tumor measurements were conducted thrice weekly using digital calipers, calculated as (*length* × *width*^2^)/2, and BLI measurements were acquired weekly. Compared to the YUMM3.2 parental cohort, no significant differences in overall survival were observed in the AKT1^E17K^, FAK^WT^, FAK^NDEL^, or FAK^Y397E^ cohorts (Figure 4A). However, mice bearing FAK^Y397E^-driven tumors exhibited prolonged survival when compared to the AKT1^E17K^ and FAK^NDEL^ cohorts, potentially due to slower primary tumor growth (Figure 4A-B). Notably, mice injected with YUMM3.2 FAK^K454R^ cells demonstrated the longest survival, and this difference was significant when compared to all other groups (Figure 4A). Additionally, the FAK^K454R^ cohort exhibited the slowest tumor growth, growing at rates slower than tumors from the parental cell line (Figure 4B). In some instances, tumors expressing FAK^K454R^ regressed completely (Figure 4C & Supplemental Figure 3). These findings suggest that the FAK^K454R^ mutant may be acting in a dominant-negative manner and highlight an essential role for FAK kinase activity in maintaining melanoma cell growth *in vivo*.

**Figure 4:**
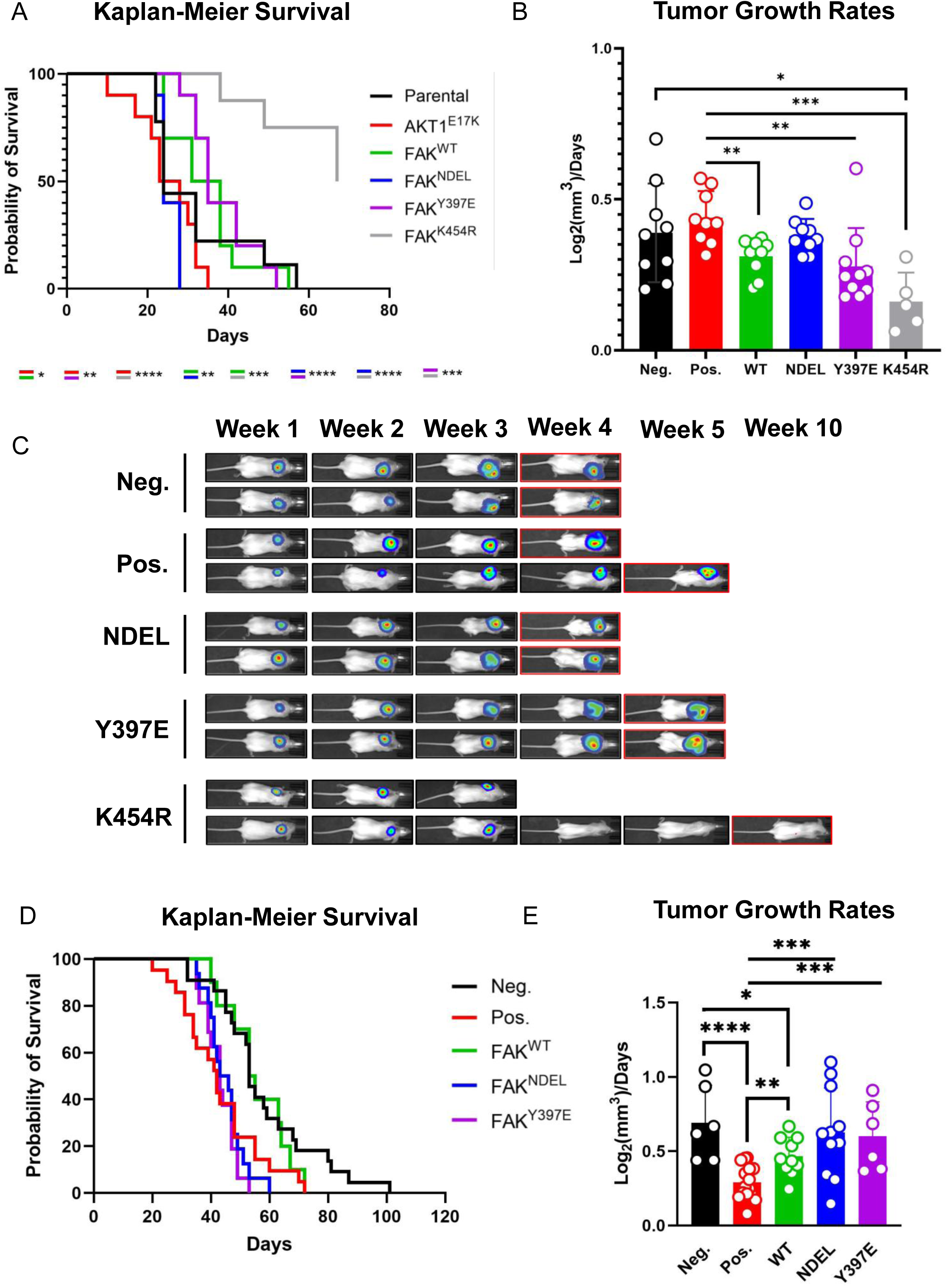
Loss of FAK activity slows tumor growth and prolongs overall survival. (A) Kaplan-Meier percent survival curves for mice injected with parental YUMM3.2 BRAF^V600E^*;Cdkn2a^−/−^;Pten^−/−^* cells (black line; negative control) or YUMM3.2 BRAF^V600E^*;Cdkn2a^−/−^;Pten^−/−^*cells expressing either AKT1^E17K^ (red line; positive control) FAK^WT^ (green line), FAK^NDEL^ (blue line), FAK^Y397E^ (purple line) or FAK^K454R^ (gray line). (B) Tumor growth rates for the mice described in A. (C) Longitudinal luminescence of the primary tumor *in vivo* as detected by BLI. (D) Kaplan-Meier percent survival curves for *Dct::TVA;Braf^CA^;Cdkn2a^lox/lox^;Pten^lox/lox^* mice injected with viruses encoding either Cre (black line; negative control), Cre + AKT1^E17K^ (positive control; red line), Cre + FAK^WT^ (green line), Cre + FAK^NDEL^ (blue line), or Cre + FAK^Y397E^ (purple line). (E) Tumor growth rates for the mice described in D. Percent brain metastasis in each cohort. P values are as follows: p < 0.05 (*), p < 0.01 (**), p < 0.001 (***), p < 0.0001 (****).

### Active FAK Reduces Survival in an Autochthonous Melanoma Mouse Model

To complement our syngeneic melanoma model, the effect of FAK^WT^ or active FAK on melanoma progression and metastasis was evaluated using the established RCAS/TVA autochthonous mouse model of melanoma. The FAK^NDEL^ and FAK^Y397E^ cDNA, along with wildtype FAK, were Gateway cloned into the RCAS vector. The proviral DNA was transfected into DF-1 avian fibroblast cells to produce infectious virus. Immunoblot analysis for the HA epitope tag confirmed the expression of all FAK constructs in DF-1 cells (Supplemental Figure 2). Newborn *DCT-TVA::Braf^CA/CA^;Cdkn2a^lox/lox^;Pten^lox/lox^*mice were subcutaneously injected with RCAS-Cre alone or in combination with RCAS-AKT1^E17K^, RCAS-FAK^WT^, RCAS-FAK^NDEL^, or RCAS-FAK^Y397E^. Post-weaning, tumor growth was monitored, and mice were euthanized when tumors reached approximately 10% of body weight or upon signs of distress. Tumor volumes were measured thrice weekly using digital calipers and calculated as (*length* × *width*^2^/2). Mice bearing tumors induced by RCAS-Cre with either RCAS-FAK^NDEL^ or RCAS-FAK^Y397E^ exhibited significantly reduced survival compared to those induced by RCAS-Cre alone, with survival outcomes comparable to those induced by the combination of RCAS-Cre and RCAS-AKT1^E17K^ (Figure 4D). Conversely, mice bearing tumors driven by FAK^WT^ did not have a significantly different overall survival compared to those driven by RCAS-Cre alone. This result mirrors our previous finding that wildtype AKT1 does not recapitulate the phenotypes of the activated AKT1^E17K^ *in vivo*^6^. Interestingly, tumor growth rates did not correlate with survival outcomes (Figures 4D&E), with tumors harboring FAK^NDEL^ and FAK^Y397E^ mutations growing at similar rates at the negative control, while tumors expressing AKT1^E17K^ and FAK^WT^ grew significantly slower (Figure 4E). These data demonstrate that in this setting, active FAK significantly shortens overall survival, although this effect is not mediated by accelerated tumor growth.

### FAK Kinase Activity Promotes Melanoma Metastasis *In Vivo*

To evaluate the effect of FAK activity on metastasis, we used both the syngeneic and autochthonous models described above. In the syngeneic model, *ex vivo* BLI on the brain, liver, and lungs was performed at euthanasia (Figure 5A-D & Supplemental Figure 3). No metastases were detected in the brain or liver of the YUMM3.2 parental cohort, and only ∼10% of mice developed lung metastases (Figure 5A-D). In contrast, ∼50% of mice with FAK^NDEL^-expressing tumors and 100% of those with FAK^Y397E^-expressing tumors developed brain metastases (Figure 5A). While not significantly increased in the other cohorts compared to the negative control, the incidence of lung and liver metastasis was also significantly increased in the FAK^Y397E^ cohort (Figure 5B-C). Notably, no metastases were observed in the brain, liver, or lung in mice injected with YUMM3.2 FAK^K454R^ cells (Figure 5A-D).

**Figure 5:**
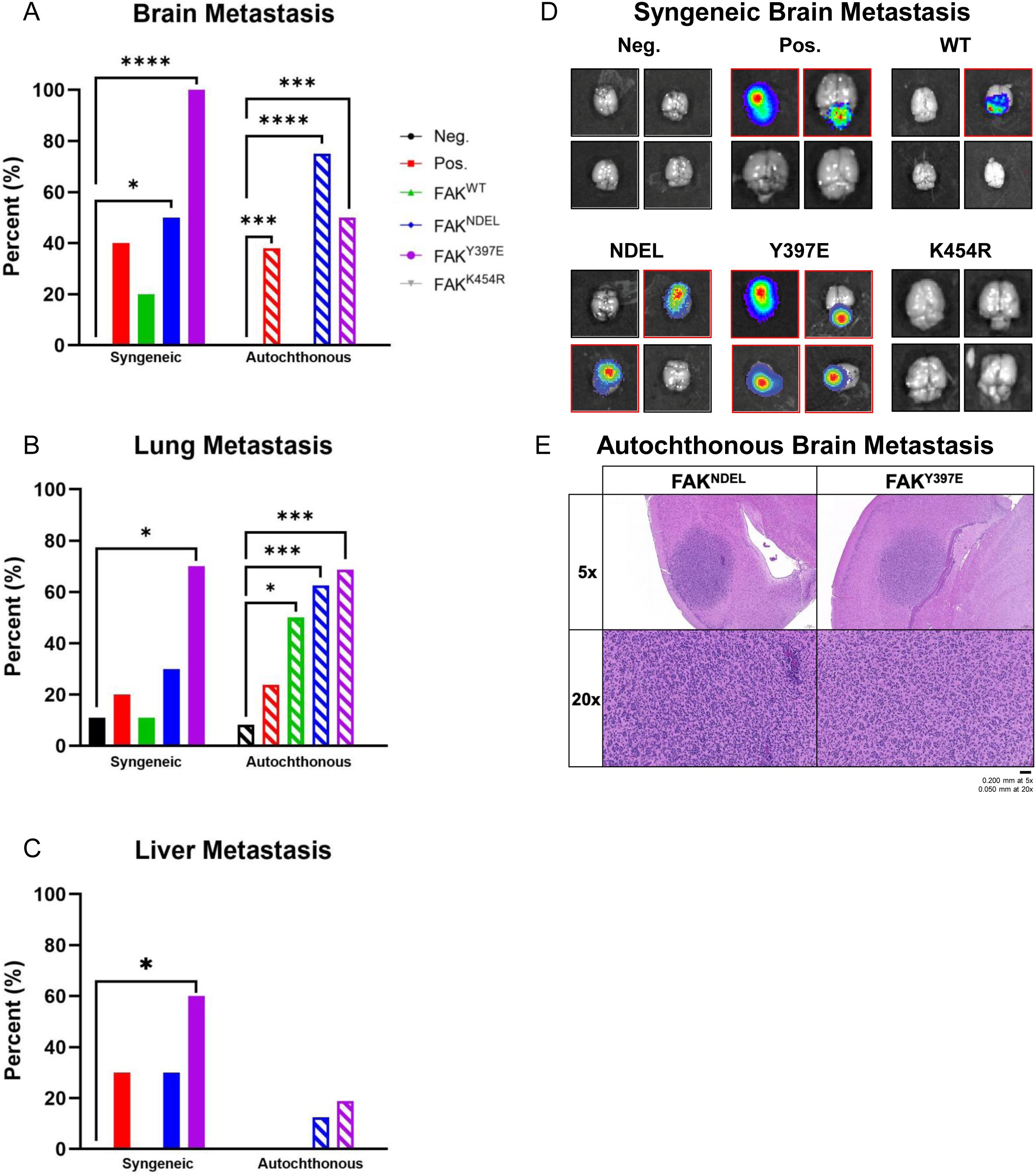
FAK kinase activity promotes melanoma metastasis. (A) Brain metastasis incidence observed in syngeneic and autochthonous models. (B) Lung metastasis incidence observed in syngeneic and autochthonous models. (C) Lung metastasis incidence observed in syngeneic and autochthonous models. (D) Representative *ex vivo* bioluminescent images of brains from mice in each cohort of the syngeneic mouse model. *Ex vivo* BLI of brain, liver, and lung was used to detect metastases reported in (A-C). (E) Representative hematoxylin and eosin (H&E) staining of brain metastases from FAK^NDEL^ and FAK^Y397E^ cohorts in the autochthonous mouse model. H&E of brain, liver, and lung was used to detect metastases reported in (A-C).

Metastasis incidence to the brain, liver, and lung in the autochthonous melanoma mouse model was assessed via hematoxylin and eosin (H&E) staining of formalin-fixed paraffin-embedded tissue collected at euthanasia (Figure 5A-C & 5E). Consistent with prior reports^5,6^, approximately 40% of the RCAS-AKT1^E17K^ cohort developed brain metastases, while incidences of ∼70% and ∼50% were observed in the RCAS-FAK^NDEL^ and RCAS-FAK^Y397E^ cohorts, respectively (Figure 5A). Further, lung metastasis incidence was significantly increased in the activated FAK cohorts, with lung metastases occurring in ∼60% in the RCAS-FAK^NDEL^ cohort and ∼70% in the RCAS-FAK^Y397E^ cohort compared to ∼10% in mice injected with RCAS-Cre alone (Figure 5B). Liver metastases were also increased in the RCAS-FAK^NDEL^ and RCAS-FAK^Y397E^ cohorts when compared to mice injected with RCAS-Cre alone, although this increase was not significant (Figure 5C). These data collectively demonstrate that FAK activity promotes melanoma metastasis in these models.

### FAK Functions Downstream of PTEN Lipid Phosphatase to Promote Brain Metastasis

Our previous studies demonstrate that *Pten* loss cooperates with active AKT1 to promote melanoma brain metastasis^5^. PTEN possesses dual phosphatase activity, whereby it can remove phosphate groups from both lipids and proteins. The canonical function of PTEN is to inhibit the PI3K/AKT signaling pathway by converting phosphatidylinositol (3,4,5)-triphosphate (PIP_3_) to phosphatidylinositol (4,5)-bisphosphate (PIP_2_) through its lipid phosphatase activity, but it has also been shown to act as a protein phosphatase to inactivate FAK by dephosphorylating FAK at Y397^34^. Interestingly, FAK can also phosphorylate PTEN at Y336, enhancing PTEN phosphatase activity in an apparent negative feedback loop^35^. To better understand how loss of *Pten* sustains FAK activity and promotes melanoma brain metastasis, we generated loss-of-function mutations of *Pten* to disrupt its lipid and/or protein phosphatase activity.

A lipid phosphatase-deficient mutant (PTEN^G129E^) and a protein phosphatase-deficient mutant (PTEN^Y138L^) were generated to explore the isolated roles of PTEN protein phosphatase activity and lipid phosphatase activity, respectively. Using the aforementioned syngeneic melanoma model, these loss-of-function PTEN mutants were expressed in YUMM3.2 cells with CRISPR-mediated knockout of wild-type PTEN, and expression of AKT1^E17K^. As previously discussed, AKT1^E17K^ was used as a positive control for *in vivo* brain metastases, allowing us to assess the ability of PTEN to reduce metastases to the brain. The expression of these proteins was confirmed via immunoblotting for the HA epitope tag and PTEN (Figure 6A). These modified YUMM3.2 cells were then subcutaneously injected into two cohorts of 6- to 10-week-old GH mice as described above. After evaluation of metastasis via *ex vivo* BLI, no metastases were observed in the mice injected with AKT1^E17K^PTEN^Y138L^ cells (Figure 6B & Supplemental Figure 4), whereas 40% of the mice injected with the cells expressing only AKT1^E17K^ developed brain metastases. In contrast, ∼75% of mice in the PTEN^G129E^ cohort developed brain metastases, ∼40% developed lung metastases, and ∼62% developed liver metastases (Figure 6B). These results are consistent with a dominant role for PTEN lipid phosphatase activity in suppressing metastasis, while recognizing that the PTEN^Y138L^ mutant may not fully isolate protein phosphatase function in every biological setting.

**Figure 6:**
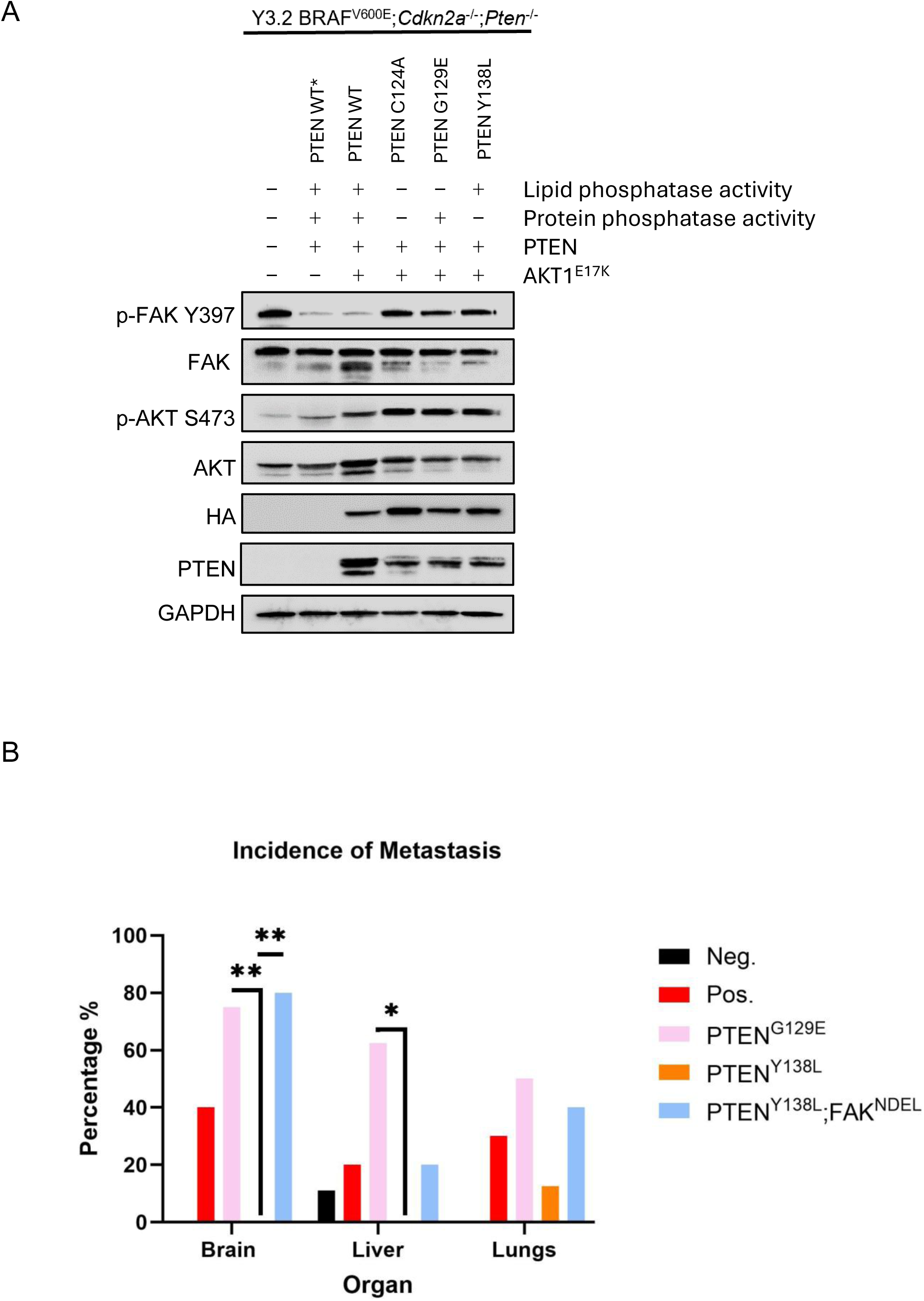
FAK overcomes PTEN lipid phosphatase activity to re-establish brain metastasis. (A) Expression of AKT1^E17K^, PTEN^WT^, PTEN^C124A^, PTEN^G129E^, and PTEN^Y138L^ was confirmed by anti-HA in YUMM3.2 BRAF^V600E^*;Cdkn2a^−/−^;Pten^−/−^* cells. Phosphorylated FAK (Y397) as well as total FAK was assessed as a measure of FAK activity. Phosphorylated AKT (S473) as well as total AKT was assessed as a measure of AKT activity. PTEN^WT*^ denotes endogenous PTEN. PTEN^C214A^, a dual lipid and protein phosphatase mutant, was used as a negative control for phosphatase activity. GAPDH was included as a loading control. (B) Brain, lung, and liver metastasis in the syngeneic model in mice injected with YUMM3.2 BRAF^V600E^*;Cdkn2a^−/−^;Pten^−/−^*cells expressing AKT1^E17K^ with the designated PTEN or FAK mutants. PTEN lipid phosphatase (AKT1^E17K^PTEN^Y138L^) completely blocks brain metastasis in the syngeneic model, which is rescued by expression of FAK^NDEL^ (AKT1^E17K^PTEN^Y138L^FAK^NDEL^) as detected by *ex vivo* BLI.

To determine if FAK expression could rescue the metastatic phenotype suppressed by PTEN lipid phosphatase activity, FAK^NDEL^ was co-expressed in the AKT1^E17K^PTEN^Y138L^ cell line and subcutaneously injected into syngeneic GH mice. FAK^NDEL^ was specifically selected because it significantly promotes metastasis (Figure 5A-C) but does not possess the N-terminal FERM domain, which may be regulated by phospholipids that are affected by PTEN lipid phosphatase activity. In this cohort of mice, we observed an ∼80% incidence of brain metastasis (Figure 6B). These data demonstrate that FAK^NDEL^ promotes metastasis even in cells that retain PTEN lipid phosphatase activity, suggesting that FAK functions downstream of PTEN to promote brain metastasis.

## Discussion

In this study, we combined analysis of a clinico-genomic dataset with two complementary mouse models to demonstrate that high FAK expression is associated with worse patient outcomes and that FAK functions as a key driver of melanoma metastasis in this context. Although FAK canonically signals at focal adhesions to regulate cell migration and invasion, its specific role in melanoma metastasis has remained poorly defined. Moreover, several non-canonical functions of FAK have been reported, involving both kinase-dependent and kinase-independent activities. Our data clarify that FAK kinase activity is required for melanoma cell survival, growth, and metastasis *in vivo*, providing strong rationale for the use of ATP-competitive FAK inhibitors in this disease.

Our *in vivo* phenotypic analyses further underscore the importance of FAK kinase activity. Tumors expressing the constitutively active FAK^Y397E^ mutant exhibited slower primary tumor growth and prolonged survival compared to the AKT1^E17K^ positive control, yet displayed a highly aggressive metastatic phenotype, with 100% incidence of brain metastasis in the syngeneic model. Consistent with our *in vitro* data, these cells appear biased toward a migratory rather than proliferative state, which may explain the reduced primary tumor growth as cells preferentially disseminate to distant sites. In contrast, expression of kinase-dead FAK^K454R^ resulted in the slowest tumor growth, with occasional tumor regression, markedly reduced metastasis, and prolonged overall survival even when compared to the parental control. Together with our *in vitro* findings that FAK^K454R^ fails to promote migration or invasion, these results indicate that FAK kinase activity is required not only for metastatic dissemination but also for efficient tumor cell survival and growth *in vivo*. This is consistent with prior work showing that pharmacological FAK inhibition with VS-4718 prolongs survival, slows tumor growth, and reduces brain metastases in melanoma models^36^.

Interestingly, deletion of the N-terminal FERM domain (FAK^NDEL^) reduced cell proliferation *in vitro*. Deletion of the N-terminal FERM domain may alter FAK subcellular localization, dimerization dynamics, or phosphorylation turnover. The FERM domain plays a key role in membrane recruitment and integrin-mediated activation, and its deletion may disrupt spatial regulation of FAK, despite preserved or enhanced catalytic potential. Although loss of the N-terminal FERM domain reduced proliferation *in vitro*, this effect did not translate into reduced tumor growth *in vivo*. This discrepancy likely reflects the complexity of the tumor microenvironment, which includes stromal and immune cells, extracellular matrix, vasculature, and soluble factors. In multiple cancer types, FAK has been shown to promote a tumor-supportive microenvironment by inducing checkpoint ligand expression^37–40^, regulating chemokine and cytokine production^41,42^, and promoting fibrosis^17^. Consistent with this concept, combination strategies incorporating FAK kinase inhibition with immune checkpoint blockade and RAF/MEK inhibition have produced significant tumor regression in syngeneic melanoma models^43^. Thus, activated FAK mutants may support tumor growth *in vivo* in part by shaping a permissive microenvironment, even when intrinsic proliferative effects are modest. Defining the specific contributions of melanoma cell-intrinsic FAK signaling versus microenvironmental remodeling remains an important area for future investigation.

At the domain level, our data further refine how FAK signals in melanoma. The FAK^NDEL^ mutant promoted migration, invasion, and metastasis, indicating that the FERM domain is dispensable for these phenotypes. This has important therapeutic implications, as it suggests that allosteric inhibitors targeting the FERM domain alone may be insufficient to block tumor growth or metastatic spread in this setting. While we did not directly test the role of the C-terminal FAT domain, prior studies have reported mixed effects of FAT deletion on FAK activity, ranging from minimal impact on kinase function^28^ to reduced activity due to impaired focal adhesion localization and dimerization^44^. As alternative FAK-targeting strategies, like allosteric inhibitors and PROTACs, continue to be developed^45^, understanding how individual domains contribute to specific biological outputs will be increasingly important.

A central mechanistic insight from our work is that FAK acts downstream of, and in opposition to, the tumor suppressor PTEN. We show that PTEN lipid phosphatase activity completely prevents FAK-mediated melanoma brain metastasis, underscoring the potency of PTEN as a barrier to metastatic progression. Strikingly, activation of FAK downstream of PTEN fully rescues the brain metastatic phenotype, positioning FAK as a critical downstream effector capable of overriding the protective effects of PTEN. These findings suggest that elevated FAK signaling may confer metastatic risk even in tumors that retain PTEN function.

From a clinical perspective, our findings carry significant prognostic and therapeutic implications. Analysis of patient datasets revealed that high FAK expression correlates with shorter overall survival in melanoma, reinforcing the clinical relevance of our mechanistic data and suggesting that FAK expression may serve as a biomarker to identify patients at higher risk for metastatic disease. Moreover, our demonstration that kinase activity is the essential mediator of tumor growth and metastasis supports the clinical development of ATP-competitive FAK inhibitors, particularly for patients with brain metastases or PTEN pathway alterations. An ongoing clinical trial is evaluating the combination of avutometinib (RAF/MEK clamp) and defactinib (FAK inhibitor), with or without encorafenib, in patients with advanced melanoma and brain metastases (NCT06194929). While early, this study will provide important insight into whether FAK inhibition can improve outcomes in this high-risk population. Given the limited efficacy of current therapies against brain metastases, targeting FAK represents a promising strategy that directly interferes with the molecular machinery driving metastatic colonization.

### Limitations to the study

Our results support the role of FAK catalytic activity in promoting melanoma metastasis. However, we did not directly test the effect of FAK on the individual steps of the metastatic cascade. Therefore, it is unclear whether this increase in metastasis incidence is primarily attributable to enhanced tumor cell survival in circulation, increased efficiency of intravasation or extravasation across the vasculature and blood-brain barrier, or improved adaptation to and survival within the microenvironment of metastatic sites. Notably, our parental cohort in the syngeneic model demonstrates that primary tumor growth rate does not correlate with metastatic potential. Parental tumors grew more rapidly than FAK^Y397E^-expressing tumors, yet the latter displayed a higher incidence of metastasis. This dissociation suggests that enhanced metastatic potential is not simply a function of proliferative capacity at the primary site. Interestingly, prior studies using pharmacologic FAK inhibition in models of established brain metastasis have demonstrated a reduction in intracranial metastatic burden, supporting a role for FAK in promoting tumor cell survival in the brain metastatic niche^36^. Whether FAK activity increases brain metastatic incidence by augmenting the number of viable circulating tumor cells that ultimately seed the brain vasculature remains unresolved.

Another limitation relates to our experimental strategy of expressing exogenous FAK mutants in cell lines that retain endogenous wild-type FAK expression. Although the kinase-dead FAK^K454R^ mutant appears to function in a dominant-negative manner, we were unable to fully ablate endogenous FAK activity. Attempts to generate CRISPR-mediated knockout clones were unsuccessful in the melanoma cell lines utilized in this study, potentially reflecting negative selection against complete FAK loss due to its roles in tumor cell survival and proliferation. CRISPR-mediated FAK knockout has been reported in the BRAF-mutant human melanoma cell line A375, where FAK loss in combination with BRAF/MEK inhibition enhanced apoptosis and reduced colony formation^43^. However, the independent effects of FAK depletion alone on cell proliferation and survival were not reported.

## Conclusions

In summary, our study provides a mechanistic framework linking FAK kinase activity to melanoma metastasis, delineates its relationship to PTEN signaling, and underscores its prognostic and therapeutic significance. These findings support the continued development and clinical testing of FAK inhibitors, with particular attention to patient stratification based on FAK expression, as a strategy to improve outcomes in metastatic melanoma.

## Declarations

### Author contributions

K.S. and S.H. designed the experiments. K.S., M.F., A.P., G.P., P.M., T.T., and D.K. performed the experiments. K.S., M.F., A.P., G.P., P.M., and S.H. analyzed the data. K.S., M.F., A.P., and S.H. wrote the manuscript. All authors discussed the results and reviewed and revised the manuscript accordingly.

## Acknowledgements

We thank members of the VanBrocklin, McMahon, and Welm labs for providing mouse strains, vectors, and/or advice. We thank the Huntsman Cancer Institute Vivarium staff for assistance with mouse husbandry. We thank Caris Life Sciences for use of their clinico-genomic database. Research reported in this publication utilized Histology, DNA Synthesis, and DNA Sequencing cores at Huntsman Cancer Institute at the University of Utah. These core facilities and shared resources are supported by the National Cancer Institute of the National Institutes of Health (NIH) under award number P30CA042014.

---

**Table 1 title:**

FAK expression correlates with worse outcomes in melanoma patients.

**Table 1 legend:**

Statistics of cohorts represented in Figure 1. TPM: transcripts per million; CI: confidence interval; HR: hazard ratio.

