## Supplementary figures and images for "Focal adhesion kinase promotes metastasis in BRAF-mutant melanoma"

### Supplemental Figure 1

#### Slide 1
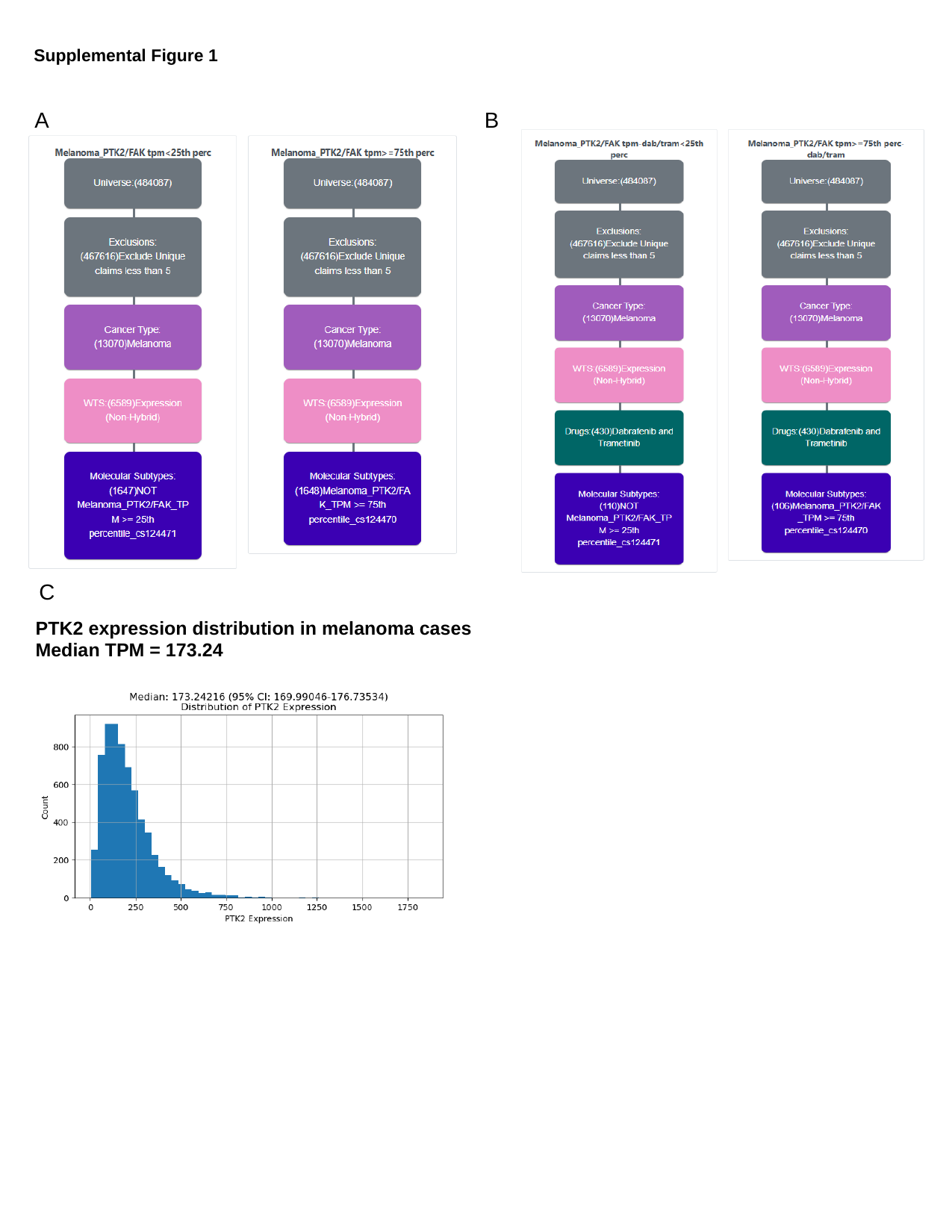

Supplemental Figure 1
A B
C
PTK2 expression distribution in melanoma cases
Median TPM = 173.24

### Supplemental Figure 3

#### Slide 1
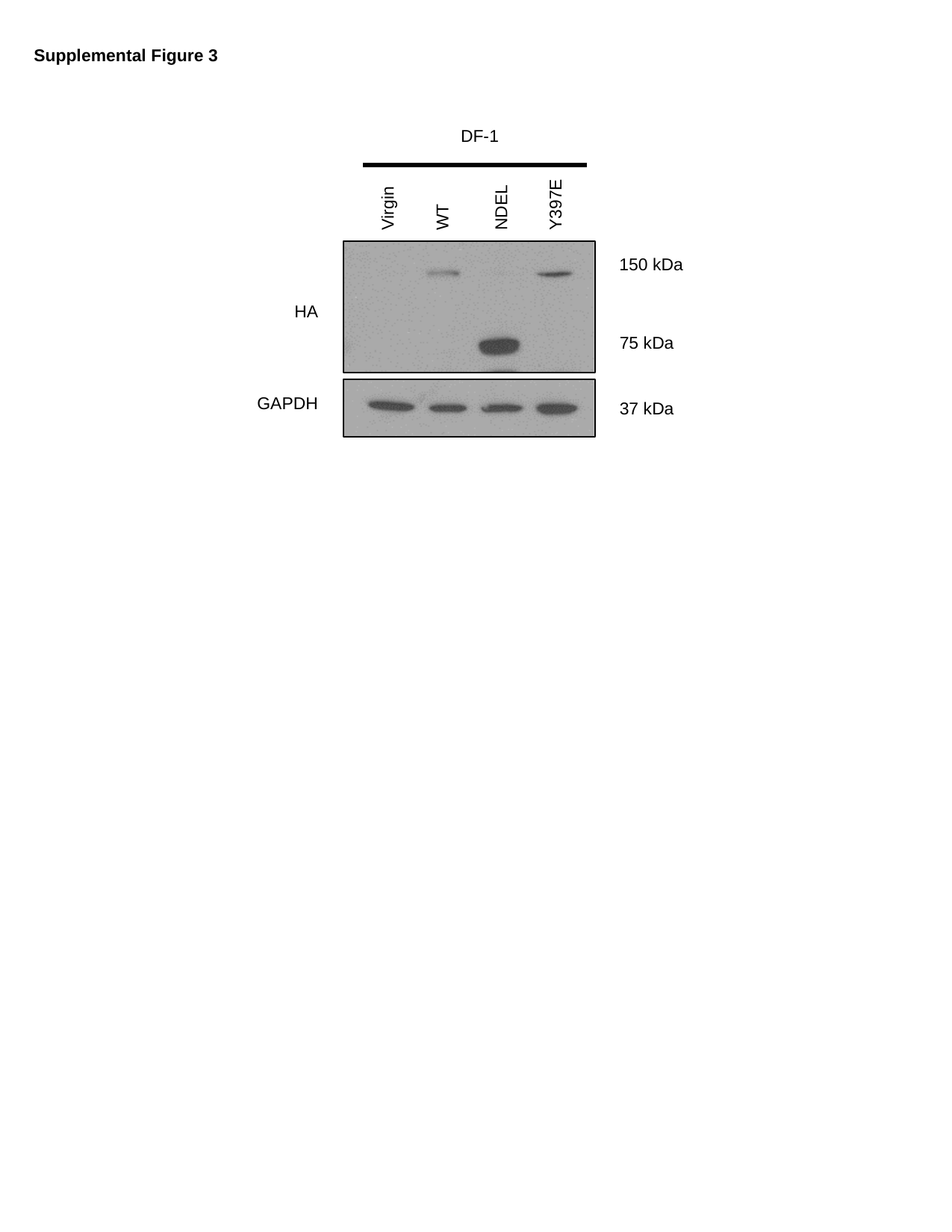

Supplemental Figure 3
DF-1
Y397E
NDEL
Virgin
WT
150 kDa
HA
75 kDa
GAPDH
37 kDa
