## Supplemental Figure 2 for "Focal adhesion kinase promotes metastasis in BRAF-mutant melanoma"

### Slide 1
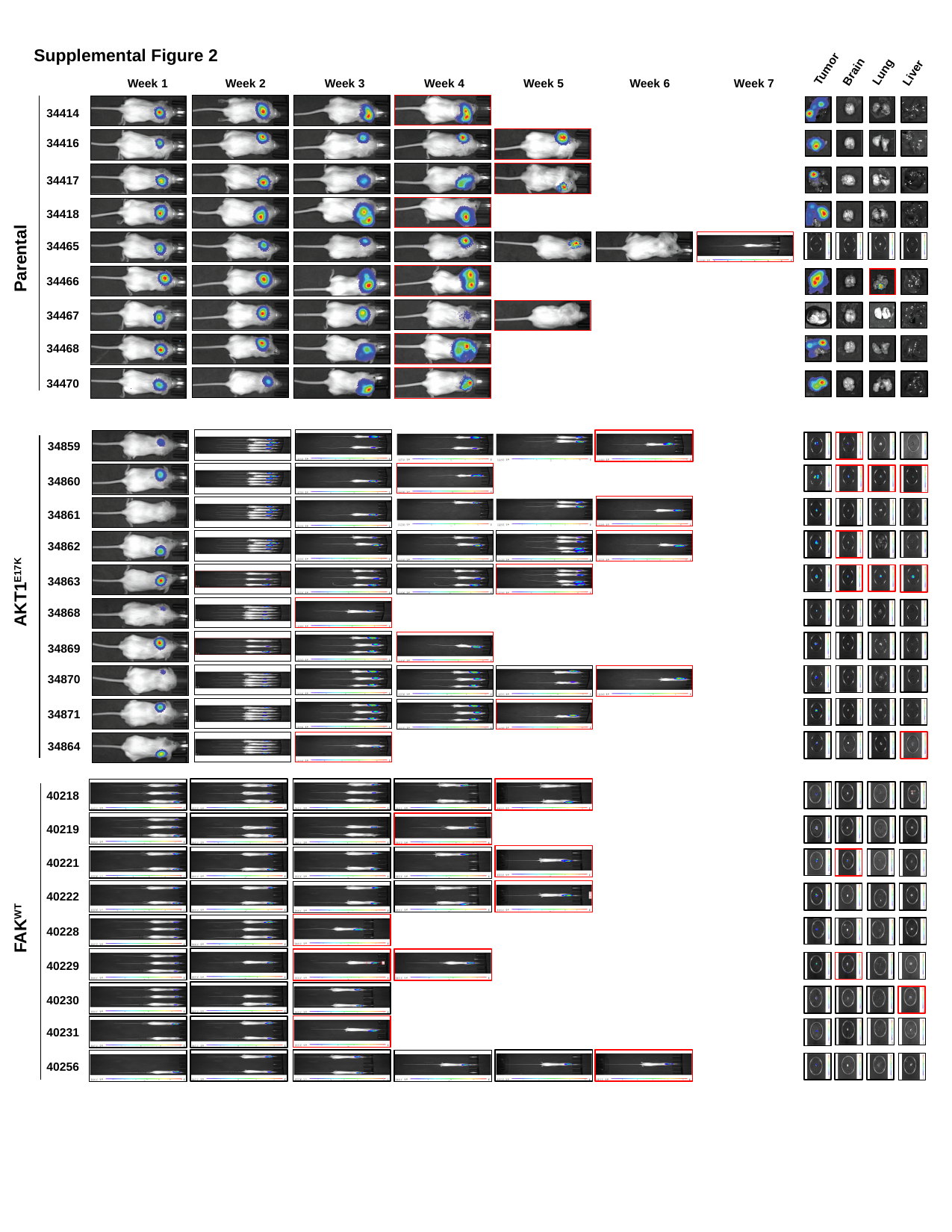

Supplemental Figure 2
Tumor
Brain
Lung
Liver
Week 1
Week 2
Week 3
Week 4
Week 5
Week 6
Week 7
34414
34416
34417
34418
34465
Parental
34466
34467
34468
34470
34859
34860
34861
34862
34863
AKT1^E17K^
34868
34869
34870
34871
34864
40218
40219
40221
40222
FAK^WT^
40228
40229
40230
40231
40256

### Slide 2
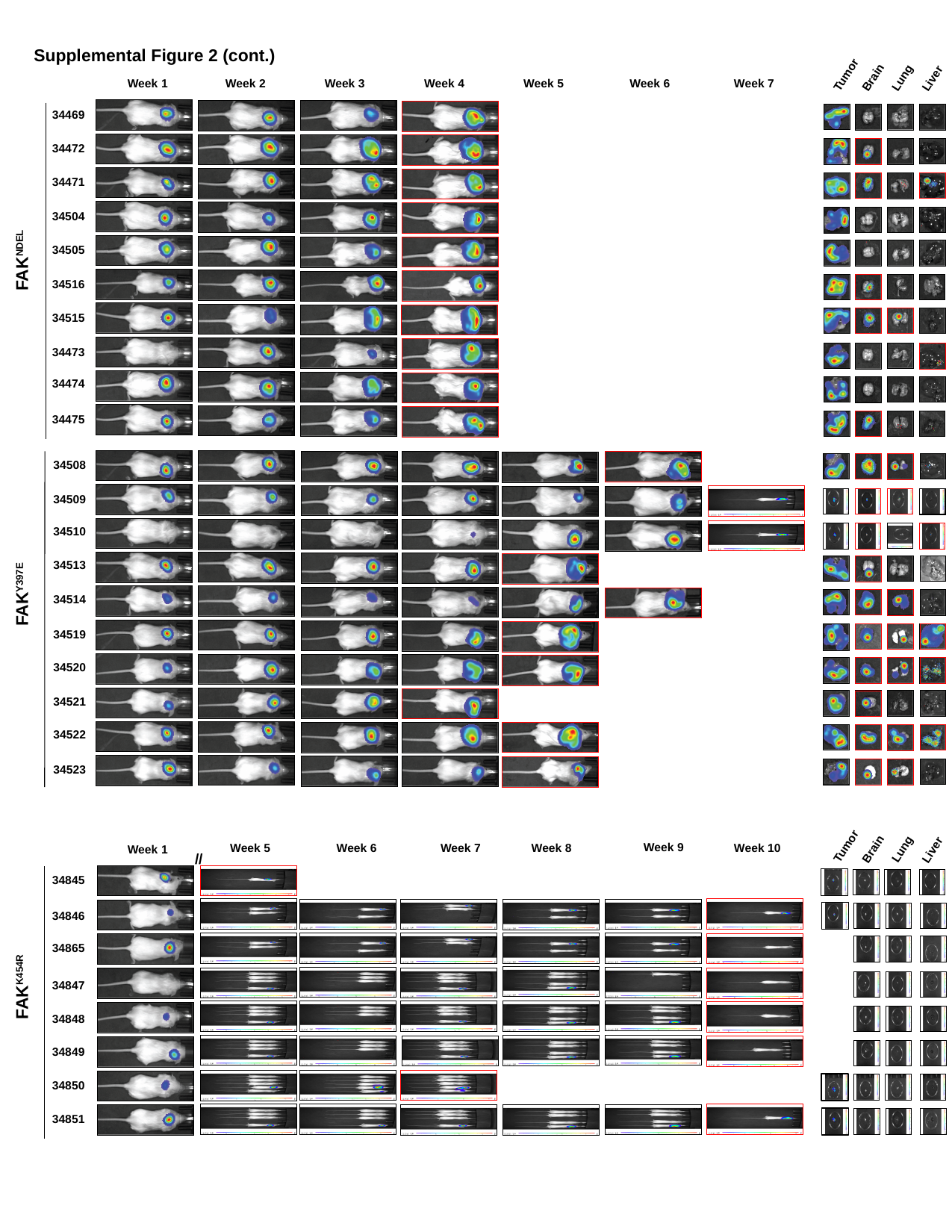

Supplemental Figure 2 (cont.)
Tumor
Brain
Lung
Liver
Week 1
Week 2
Week 3
Week 4
Week 5
Week 6
Week 7
34469
34472
34471
34504
34505
FAK^NDEL^
34516
34515
34473
34474
34475
34508
34509
34510
34513
34514
34519
34520
34521
34522
34523
FAK^Y397E^
Tumor
Week 9
Week 5
Week 6
Week 7
Week 8
Week 10
Brain
Lung
Week 1
Liver
^//^
34845
34846
34865
34847
34848
34849
34850
34851
FAK^K454R^
