## Supplemental Figure 4 for "Focal adhesion kinase promotes metastasis in BRAF-mutant melanoma"

### Slide 1
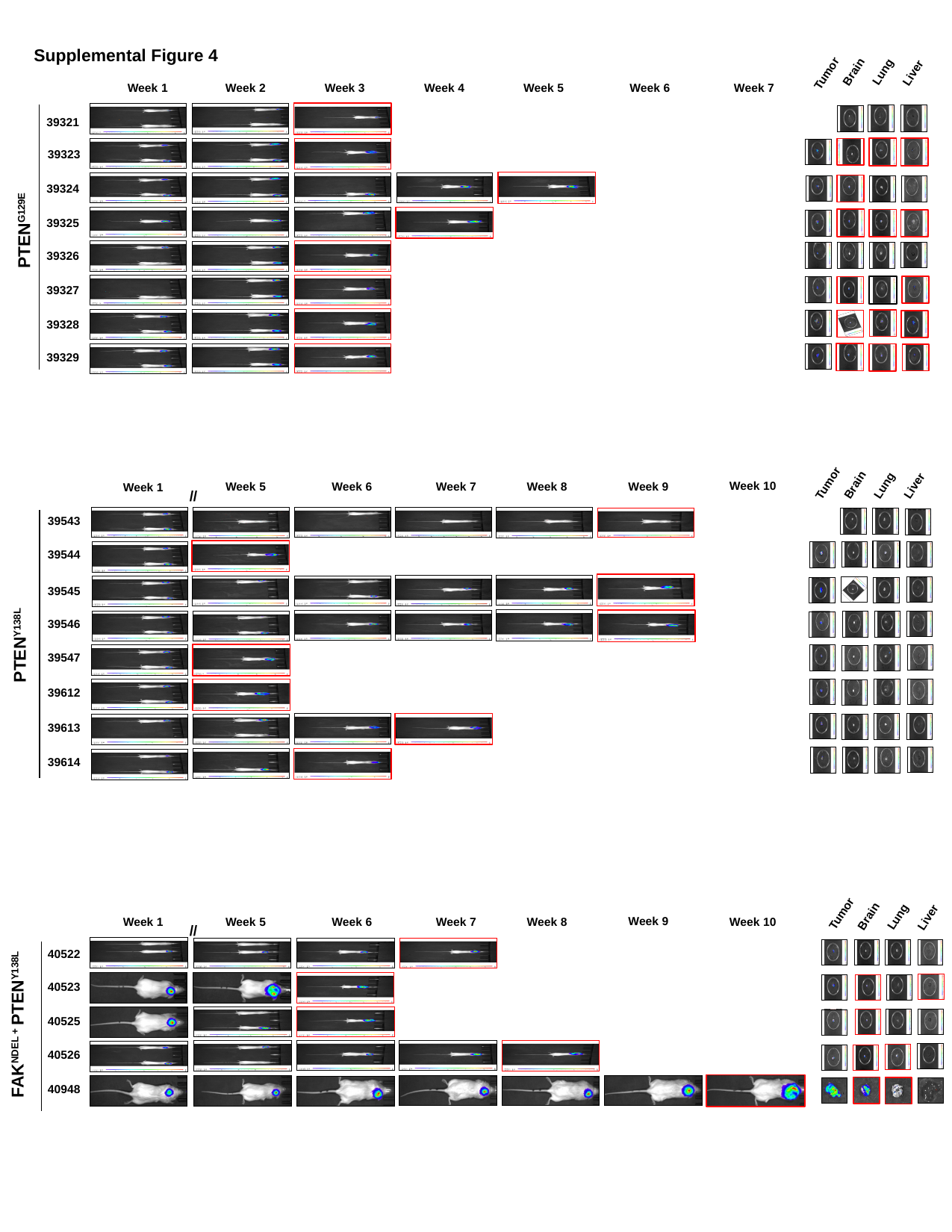

Supplemental Figure 4
Brain
Lung
Liver
Tumor
Week 1
Week 2
Week 3
Week 4
Week 5
Week 6
Week 7
39321
39323
39324
39325
PTEN^G129E^
39326
39327
39328
39329
Tumor
Brain
Lung
Liver
Week 10
Week 9
Week 5
Week 6
Week 7
Week 8
Week 1
^//^
39543
39544
39545
39546
PTEN^Y138L^
39547
39612
39613
39614
Tumor
Brain
Lung
Liver
Week 9
Week 5
Week 6
Week 7
Week 8
Week 1
Week 10
^//^
40522
40523
40525
FAK^NDEL +^ PTEN^Y138L^
40526
40948
