## Supplemental Table 1 for "Focal adhesion kinase promotes metastasis in BRAF-mutant melanoma"

### Slide 1
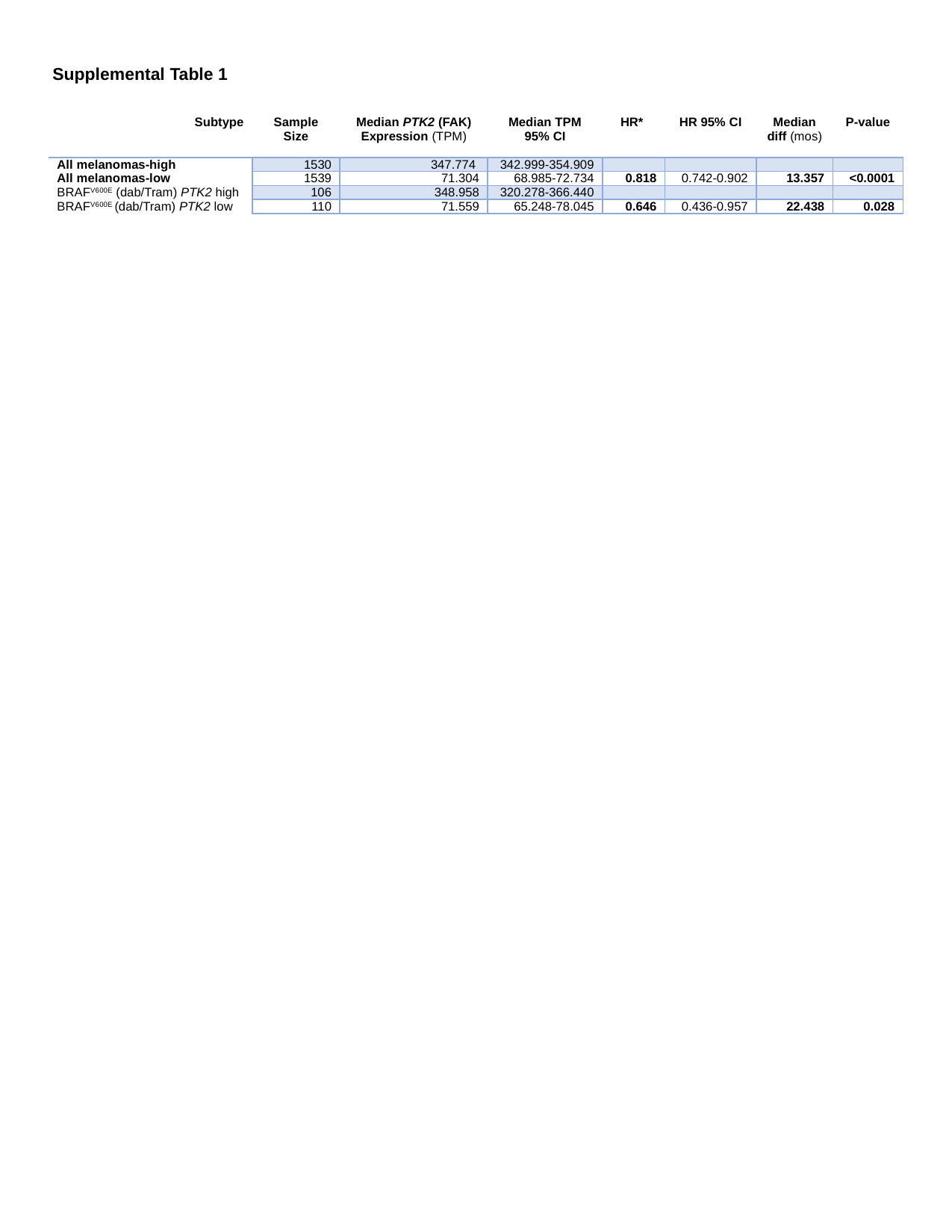

Supplemental Table 1
| Subtype | Sample Size | Median PTK2 (FAK) Expression (TPM) | Median TPM 95% CI | HR\* | HR 95% CI | Median diff (mos) | P-value |
| --- | --- | --- | --- | --- | --- | --- | --- |
| All melanomas-high | 1530 | 347.774 | 342.999-354.909 | | | | |
| All melanomas-low | 1539 | 71.304 | 68.985-72.734 | 0.818 | 0.742-0.902 | 13.357 | <0.0001 |
| BRAF^V600E^ (dab/Tram) PTK2 high | 106 | 348.958 | 320.278-366.440 | | | | |
| BRAF^V600E^ (dab/Tram) PTK2 low | 110 | 71.559 | 65.248-78.045 | 0.646 | 0.436-0.957 | 22.438 | 0.028 |
